# Balancing Growth: GROOT Genes Link Plant Biomass and Temperature Adaptation in Arabidopsis

**DOI:** 10.1101/2024.09.25.615018

**Authors:** Samsad Razzaque, Ling Zhang, Ana Paez-Garcia, Yosemite Caputi, Wolfgang Busch

## Abstract

Root systems take up water and nutrients from the soil and thereby underpin all essential plant functions. The size of the root system affects the ability of roots to capture these resources and determine the amount of carbon that roots transfer into the soil. Understanding the genetic basis of root biomass is therefore very important for enhancing crop resilience and productivity, especially in the face of climate change, as for soil carbon sequestration efforts. In our study, we catalogued root biomass of a diverse set of Arabidopsis accessions mainly derived from Sweden and Spain and utilized GWAS to identify loci associated with root biomass in *Arabidopsis thaliana*. We found that genetic variants associated with high biomass are enriched in accessions originating from distinct biogeographic regions in Spain. Among the most significant SNPs, one SNP on chromosome 3, was closely linked to three genes for which loss of function mutations caused significant increases in root, shoot and seed biomass and that we named *GROOT* genes. These genes act as general growth limiters. Additionally, our analysis revealed that the *GROOT* genes are in strong linkage disequilibrium, implying a potential coordinated function in regulating growth. Furthermore, accessions carrying the non-reference allele at this SNP showed markedly higher biomass under elevated temperatures, suggesting that these genes may also play a role in temperature adaptation. The discovery that the *GROOT* genes can enhance biomass without negative trade-offs across multiple traits opens new avenues for further research aimed at understanding genetically determined growth limitations, and for improving crop resilience and adaptability, as well as carbon sequestration.

## Introduction

Roots are essential for most land plants as they forage the soil for nutrients and water, which are critical for their growth and survival and provide anchorage for the plant body (Ogura et al. 2019; Maurel and Nacry 2020). Root systems are important for plant resilience, particularly in the face of climate change and temperature fluctuations (Mayjonade et al. 2019; Hancock et al. 2011; Karlova et al. 2021). Given the ever-changing environment, with significant fluctuations in various environmental variables, roots exhibit remarkable plasticity in their growth patterns, allowing plants to adapt and thrive in their natural habitats (Lorts and Lasky 2020; Fitz Gerald et al. 2006; Gruber et al. 2013). The extensiveness of the root system is determined by its growth rate. The cumulative growth of a root system can be measured as root biomass. Larger root systems promise to contribute significantly to carbon sequestration, as they store carbon captured through photosynthesis, with deeper, more extensive root systems enhancing soil carbon storage (Kumar et al. 2006; Panchal et al. 2022). Increased soil inputs via larger root systems also enhance soil health and fertility, fostering sustainable ecosystems (Oren et al. 2001; Sang et al. 2013).

Root biomass is a complex trait likely influenced by pleiotropic effects involving multiple genes and genetic pathways (Chen et al. 2021). While not well understood, the genetic basis for root biomass accumulation is shaped by various factors, including resource allocation strategies, which can create trade-offs between root and shoot growth (Lynch 2022). For instance, plants that invest more in root development might experience reduced above-ground growth. However, these trade-offs are not always straightforward and can be context dependent. Environmental conditions, such as soil fertility, water availability, and light intensity, can influence the extent and nature of these trade-offs. In nutrient-rich environments, plants may be able to support both robust root and shoot growth, whereas in nutrient-poor conditions, prioritizing root growth for better resource acquisition may come at the expense of shoot development. Moreover, environmental fluctuations and community assembly can alter this dynamic, as different conditions and community compositions may select for varying root and shoot traits (Lavorel and Garnier 2002; García-Palacios et al. 2013; Gedroc et al. 1996). Additionally, genetic factors and the specific roles of different genes can modulate these trade-offs, suggesting that the interplay between root and shoot growth is highly dynamic and influenced by a myriad of internal and external factors (He et al. 2022; Dwivedi et al. 2021; Smakowska et al. 2016; Lundgren and Des Marais 2020).

Only a few genes are known to simultaneously enhance both shoot and root biomass. Overexpression of the *CYTOKININ OXIDASE (CKX)* gene generally accelerates and enlarges root growth but significantly reduces leaf production to just 3-4% of that in wild-type plants (Werner et al. 2001). This reduction may be due to *CKX* overexpression in both root and shoot tissues. However, tissue-specific expression of the *CKX* gene can modulate the trade-off between root and shoot growth. For example, transgenic maize with root-specific *CKX* expression showed up to 46% more root dry weight without affecting shoot growth (Ramireddy et al. 2021). Similarly, another study found that overexpressing the chickpea *CKX* gene (*CaCKX6*) under a root-specific promoter in both Arabidopsis and chickpea significantly increased lateral root number and root biomass, while leaving shoot growth unaffected (Khandal et al. 2020). A similar pattern of balancing shoot and root growth has also been observed in other genes, such as *OsWOX11* in rice and *TaNAC69-1* in wheat. *OsWOX11* enhances root biomass by promoting the growth of crown roots without affecting shoot growth (Jiang et al. 2017). In contrast, *TaNAC69-1* increases primary seminal root length and overall root biomass, as well as shoot biomass when overexpressed using specific promoters (Chen et al. 2016). The pattern of root to shoot growth under the same gene activity is inconsistent, and growth stimulation appears to be more dependent on the promoters driving the gene. Thus, it would be highly beneficial to identify genes that increase both shoot and root biomass without relying on tissue-specific promoters, as such identification could significantly impact plant performance and resilience. Therefore, identifying genes and genetic pathways that limit growth and upon loss of function enhance root and shoot growth would be highly beneficial for optimizing overall plant performance and resilience.

In this study, we explored natural variation in the root biomass of *Arabidopsis thaliana* to identify genes associated with root biomass traits using a genome wide association study (GWAS). By analyzing 52 candidate genes from the top five most significant GWAS hits, we identified three genes (*GROOT1*, *GROOT2*, *GROOT3*) that when mutated lead to significantly increased root biomass. Loss of function of these genes did not only cause a strong increase of root biomass but also influenced other traits, including increasing aboveground biomass and seed size. While being involved in distinct growth-related molecular processes, the natural alleles of the three *GROOT* genes are genetically linked to one another. Prompted by the geographic distribution of the accessions harboring distinct *GROOT* alleles, we studied the impact of the genes and alleles on growth responses to temperatures and found a significant gene and genotype by temperature interactions. Our data suggest a role of these genes and their variants for determining the overall growth and fitness of the plant and expose them as key targets for improving carbon sequestration, crop resilience and productivity in the face of climate change.

## Results

### Natural variation of root biomass in *Arabidopsis thaliana*

We set out to assess natural variation of root biomass in *Arabidopsis thaliana* (Arabidopsis) by measuring root biomass in 21-day old seedlings of Arabidopsis natural accessions. We chose 264 accessions of the 1001 genomes collection (Alonso-Blanco et al. 2016; Weigel and Mott 2009), selecting mostly accessions originating from Spain and Sweden, as Arabidopsis populations from Spain and Sweden derive from environments with diverse ecological conditions: Spain encompasses a range of climates, from Mediterranean to mountainous, offering a wide variety of habitats for Arabidopsis. Similarly, Sweden’s diverse geography, with its mix of lowland areas and alpine regions, presents unique environmental conditions. We reasoned that these distinct conditions make them valuable subjects for studying trait diversity and local adaptation. We grew twelve seedlings from each of the 264 accessions, with three seedlings on the surface of a ½ MS medium agar plate. Therefore, we had a total of four plates per accession. After 21 days of growth, we scanned each plate using the BRAT scanner system (Slovak et al. 2014). We noticed considerable differences in root growth patterns among the accessions we grew for this experiment. Figure 1a illustrates a few examples highlighting the variations in root growth patterns.

**Figure 1:**
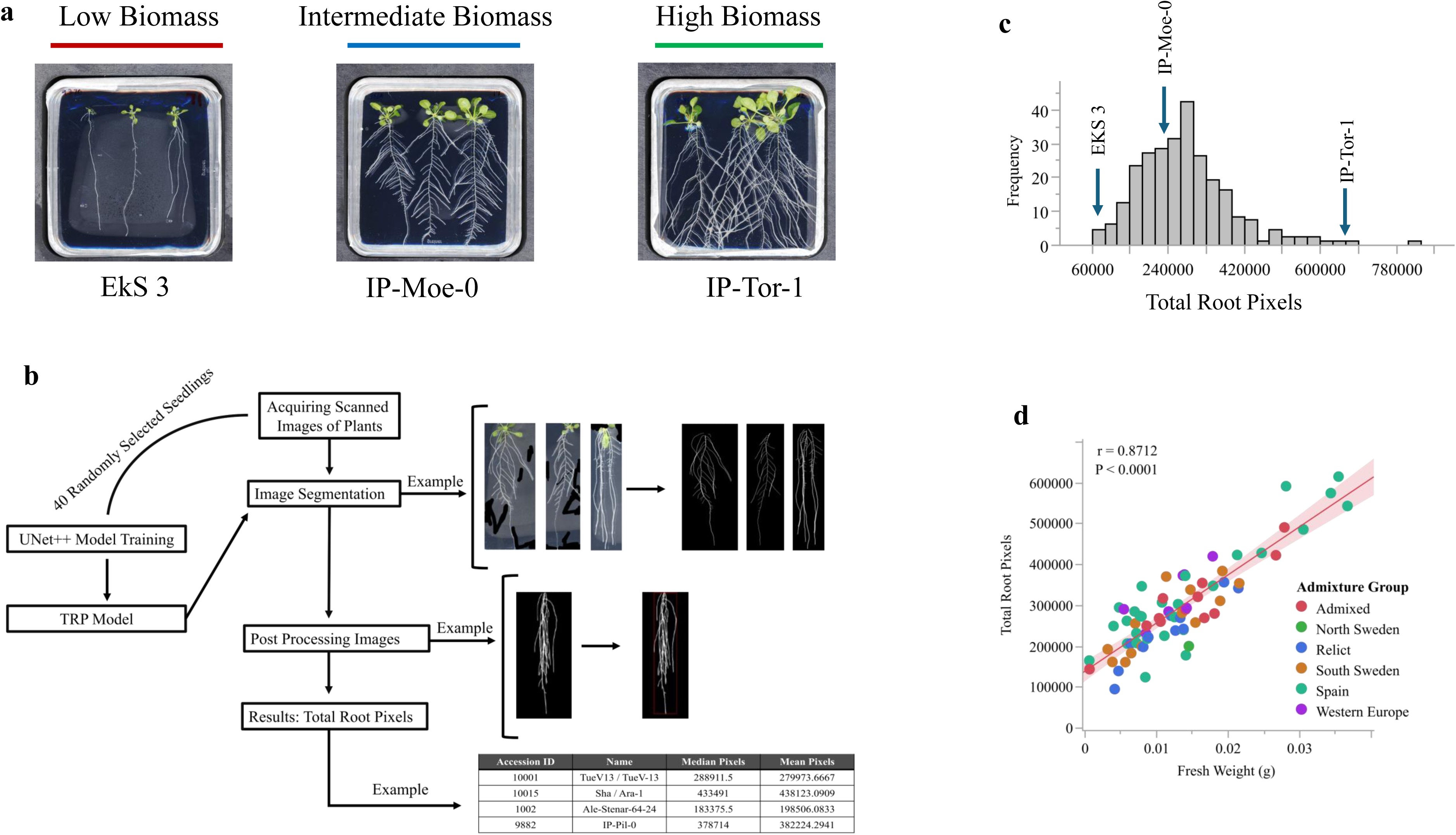
Natural variations of Arabidopsis, its quantification pipeline, and validation for root biomass trait. (a) Root growth patterns observed in Arabidopsis accessions with varying biomass production after 21 days of growth on ½ MS media under long day conditions (16h/8h). Images depict accessions with low, intermediate, and high root biomass. (b) Schematic representation of the Total Root Pixel (TRP) pipeline for estimating biomass directly from root images by tallying total pixel counts. (c) Distribution pattern of TRP across the natural accessions in this study. x-axis: total pixel number; y-axis: frequency of accessions across TRP estimates. The distribution is centered around the mean value, with three lines highlighted corresponding to the accessions depicted in (a). (d) Correlation plot between fresh weight (g) and TRP data from TRP pipeline. Each dot represents the mean root biomass value of a natural accession from a representative sample of 72 randomly selected accessions used for validation purposes. Major population groups are indicated by different colors.

As weighing single Arabidopsis roots is technically challenging and error prone due to their low mass, we decided to perform an image-based quantification of the root size. To facilitate this, we developed a deep learning UNet^++^ based pipeline to automatically quantify the number of pixels from seedlings from a scanned plate (Fig. 1b). We coined this the Total Root Pixels (TRP) pipeline (Fig. 1b) and used to quantify the number of root pixels in each of the scanned images. The distribution of average TRP of accession spanned a substantial range, from 83,938 pixels to 824,464 pixels, with an average of 278,203 pixels (Supplementary File 2). The distribution of the TRP of the natural accessions is illustrated in Fig. 1c.

We then wanted to validate that the pixel count was a good proxy for root weight. For this, we grew a randomly selected subset of the 72 natural accessions for 21 days. We then harvested the roots to measure their mass. Consistent with our assumption that TRP is a good and practical proxy measure for root fresh weight, we observed a strong positive correlation (r = 0.879, P < 0.001) between fresh weight in grams and the TRP dataset obtained using the TRP pipeline (Supplementary File 3, Fig. 1d).

Next, we wanted to test whether the proportion of total phenotypic variation of TRP among the studied lines could be attributed to genetic variation. For this, we calculated the broad-sense heritability (H²) of TRP and found it to display a high heritability of 64% (bootstrap-based significance, P < 0.001, Supplementary File 4), indicating that a majority of the observed phenotypic variance in total root pixel data is due to genetic variation. Taken together we showed that there is substantial and heritable variation for root biomass in Arabidopsis natural accessions.

### Genetic variants associated with high biomass are enriched in accessions originating from distinct biogeographic regions in Spain

To identify associations between genetic variants within the natural accessions and the root biomass, we used the TRP data to conduct a genome-wide association study (GWAS). For this, we used a linear mixed model EMMAX (Kang et al. 2010) linking the SNP data extracted from 1001 Genomes data base (full imputed; https://1001genomes.org/). We then identified significantly associated SNPs by applying a 5% false discovery rate (FDR) threshold, adjusted using the Benjamini-Hochberg (BH) procedure. This analysis revealed 43 significant associations across chromosomes (Fig. 2a, Supplementary File 5). The top five most significant hits, which not only exceeded the 5% FDR threshold but also the more stringent Bonferroni 5% threshold, were selected for candidate gene discovery. All of these hits were located on chromosomes two and three (Fig. 2a; Supplementary File 1 & 5).

**Figure 2:**
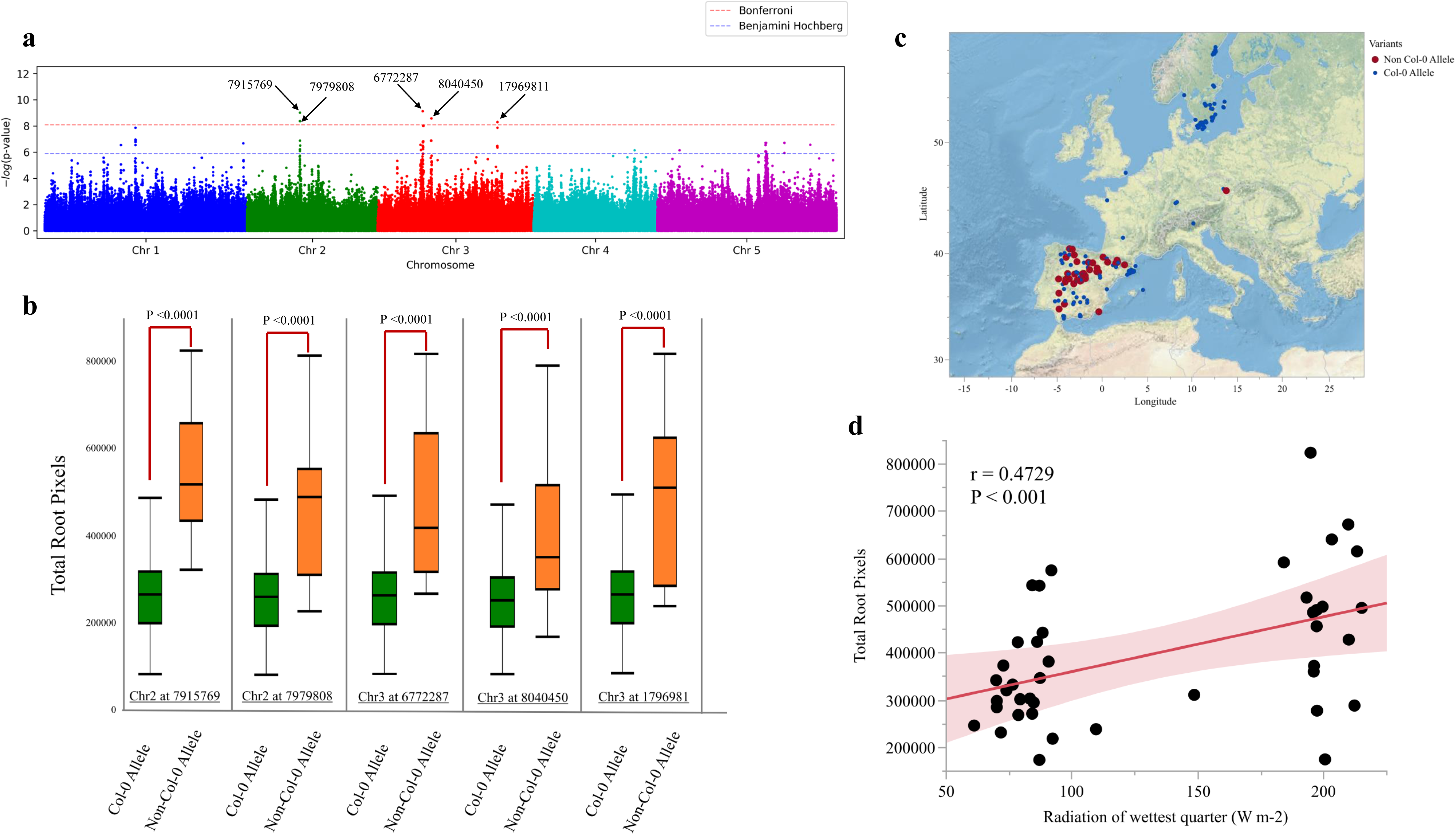
Genome-wide association study (GWAS) identifies loci associated with biomass and reveals patterns linked to local climates. (a) Manhattan plot of GWA mapping for root biomass (TRP) after 21 days of growth on ½ MS media. Horizontal blue line: 5% false discovery rate threshold; horizontal red line: Bonferroni 5% threshold. Top five associated SNPs are highlighted. Colors indicate different chromosomes (b) Box plots of the top five SNPs and their allelic variation within the natural accessions used in this study. Y-axis: TRP values; X-axis: SNPs. Non-Col-0 allele accessions exhibit significantly higher TRP values. In box plots, the horizontal line indicates the mean value, the lower and upper edges represent the 25th and 75th percentiles, and the whiskers extend to the minimum and maximum values. (c) Physical map of accession collection sites; each dot represents an accession; colors indicate SNP variants for the top five GWAS hits. (d) Correlation between TRP (y-axis) and Bio24 (Radiation of the wettest quarter in Wm-2; x-axis) for accessions exhibiting non-Col-0 allele variants for the top five SNPs. Correlation coefficients (r) and significance levels (P-values) are indicated.

The most significantly associated SNP (marker SNP) on chromosome 2 (position 7915769) captured a notable increase (103%) in pixel counts for the non-reference allele compared to the reference (Col-0) allele (P < 0.0001), as depicted in Fig. 2b. Across four other significant hits from the GWAS dataset, we consistently observed higher pixel counts associated with the non-reference allele (P < 0.05), indicating its contribution to greater pixel counts (Fig. 2b). Notably, at Chromosome 2 at position 7979808bp, the non-reference allele exhibited a 75% increase in pixel counts compared to the reference allele, while at Chromosome 3 at position 6772287bp, the SNP association showed an 83% increase (Fig. 2b). Similar trends were observed at Chromosome 3 at position 8040450bp and Chromosome 3 at position 17969811bp, with the non-reference alleles displaying 56% and 82% increases, respectively, in pixel counts compared to their reference counterparts (Fig. 2b). Collectively, these findings underscore the diverse impact of allele variation on pixel counts and emphasize the significance of these genomic regions in influencing root biomass. This observation also suggests the possibility of natural selection exerting an influence on these loci, leading to the accumulation of pixel counts associated with the non-reference allele. This intricate interplay underscores the complexity of genetic associations and hints at the adaptive potential inherent within these genetic variants.

Having identified SNPs that are significantly associated with natural variation of root biomass, we set out to explore whether there are geographical patterns of SNP distribution. We observed that nearly every non-reference allele variant for the top five SNPs, which passed the Bonferroni multiple corrections and was associated with increased root biomass, was found in accessions from Spain. (Fig 2c, Supplementary File 6). This intriguing pattern suggested that either genetic relatedness or geographic or environmental factors within this region played a role in shaping the accumulation of higher root biomass associated variants. We first tested whether relatedness explained the biomass accumulation (as opposed to environmental factors). We found that neither country of origin (P = 0.435) nor population structure (P = 0.5745) explained TRP to a significant extent while genetic variants for the top five GWAS hits demonstrated a significant association (P < 0.0001), highlighting that distinct genetic variants play important role in determining root biomass accumulation patterns.

Next, we wanted to explore potential environmental factors linked to genetic variants associated with increased root biomass accumulation in natural accessions. To achieve this, we focused our analysis on populations outside of Sweden, as the top five variants correlated with increased biomass were exclusively found in non-Swedish accessions (Fig 2c, Supplementary File 6), and individuals from Sweden exhibited distinctiveness within the dataset. Within the subgroup of non-Swedish accessions, we assessed the correlation between biomass data and bioclimatic variables (Bio01-Bio035, (Fick and Hijmans 2017; Kriticos et al. 2012). We observed no significant relationship between the biomass data and any of the bioclimatic variables within that subpopulation after applying FDR correction to the P-values obtained from statistical tests (Supplementary File 7). As non-reference allele variants were associated with increased root biomass, we then focused only on the accessions with the non-reference allele for the top five GWAS SNPs (Fig. 2a) and computed correlations between biomass and bioclimatic variables. We found one significant correlation between biomass data and Bio024 (Radiation of the wettest quarter (Wm^-2^)) (r = 0.47, P < 0.05, Fig. 2d). To investigate whether the correlation between the two traits was influenced by population structure, we performed a Principal Component Analysis (PCA) on the data (biomass and Bio24) to capture the main axes of variation that might correspond to population structure. The first two principal components (PC1 explains 73.4% of the variation, and PC2 explains 26.2%, together accounting for 99.6% of the total variation.) were extracted and included in subsequent analyses. We then fit two linear regression models to account for population structure. The first model included population as a categorical variable, while the second model included PC1 and PC2 as covariates. The results indicated that Bio24 was significantly associated with biomass after accounting for population structure, whether using population as a categorical variable (P < 0.05) or using the principal components (P < 0.05). This suggests that the observed correlation between biomass and Bio24 is not solely due to population structure.

Overall, our analysis suggests that accessions may be selectively adapted to specific climatic conditions where solar radiation during the wettest quarter is highest, favoring allele variants associated with increased root biomass accumulation in their local habitats. These findings underscore the intricate interplay between genetic variation, environmental pressures, and adaptation. It suggests that local environments, such as those found in Spain, exert a significant influence on the evolutionary trajectories of plant populations, driving the accumulation of beneficial alleles associated with increased biomass accumulation.

### Loss of function mutations of GWAS candidate genes result in a significant increase of root and shoot biomass, and seed size

Our GWAS analysis had resulted in the detection of several highly significantly associated loci for root biomass. We therefore set out to identify genes that underlie the observed variation of root biomass in proximity of the identified loci. To this end, we considered candidate genes within 4,000 base pairs of each of the top five significantly associated SNPs, as identified after Bonferroni correction (Supplementary File 1). Due to the number of candidate genes, we focused on candidate genes harboring SNPs within their coding regions and did not pursue genes with SNPs in their upstream or downstream sequences. This reduced our list to 58 genes. For each of these 58 genes, we only considered those with single T-DNA insertion lines available from the stock center, further narrowing our list to 51 candidates. For these T-DNA lines, we conducted phenotyping for root biomass after confirming homozygous mutant insertion lines, comparing biomass to the wildtype (Col-0) collected from three different sources. We identified T-DNA lines that displayed higher root biomass compared to three Col-0 wildtypes that we had obtained from different sources. While the vast majority of the screened T-DNA lines exhibited no significant differences for increased root biomass compared to the wildtype or produced inconsistent results, T-DNA lines for three distinct genes (*AT3G19440*, *AT3G19590*, *AT3G19630*) consistently displayed significantly increased root dry weight (g) compared to the three wildtype lines (Supplementary Figure 1). Loss of function mutants of each of these three genes demonstrated not only statistically significant (P < 0.05) higher mass for root mass after a 21-day growth period on MS plates, but also for shoot mass (Figure 3a). In addition to increased root and shoot biomass, we did not observe any abnormal growth or any other obvious phenotypes in plants of these lines. Taken together, our data indicate that each of these three genes has a function in limiting growth (given that loss of function mutant yielded in higher biomass). We therefore named these previously uncharacterized genes *GROOT1* to *GROOT3* (*GRT*, based on the finding that mutations of these genes lead to **G**reater **ROOT** biomass and the fictional comic book character from a large, tree-like alien species that originated from Planet X). Interestingly, all these three genes for which we had identified increased biomass upon their mutation were on chromosome 3 and originated from the GWAS peak at position 6772287.

**Figure 3:**
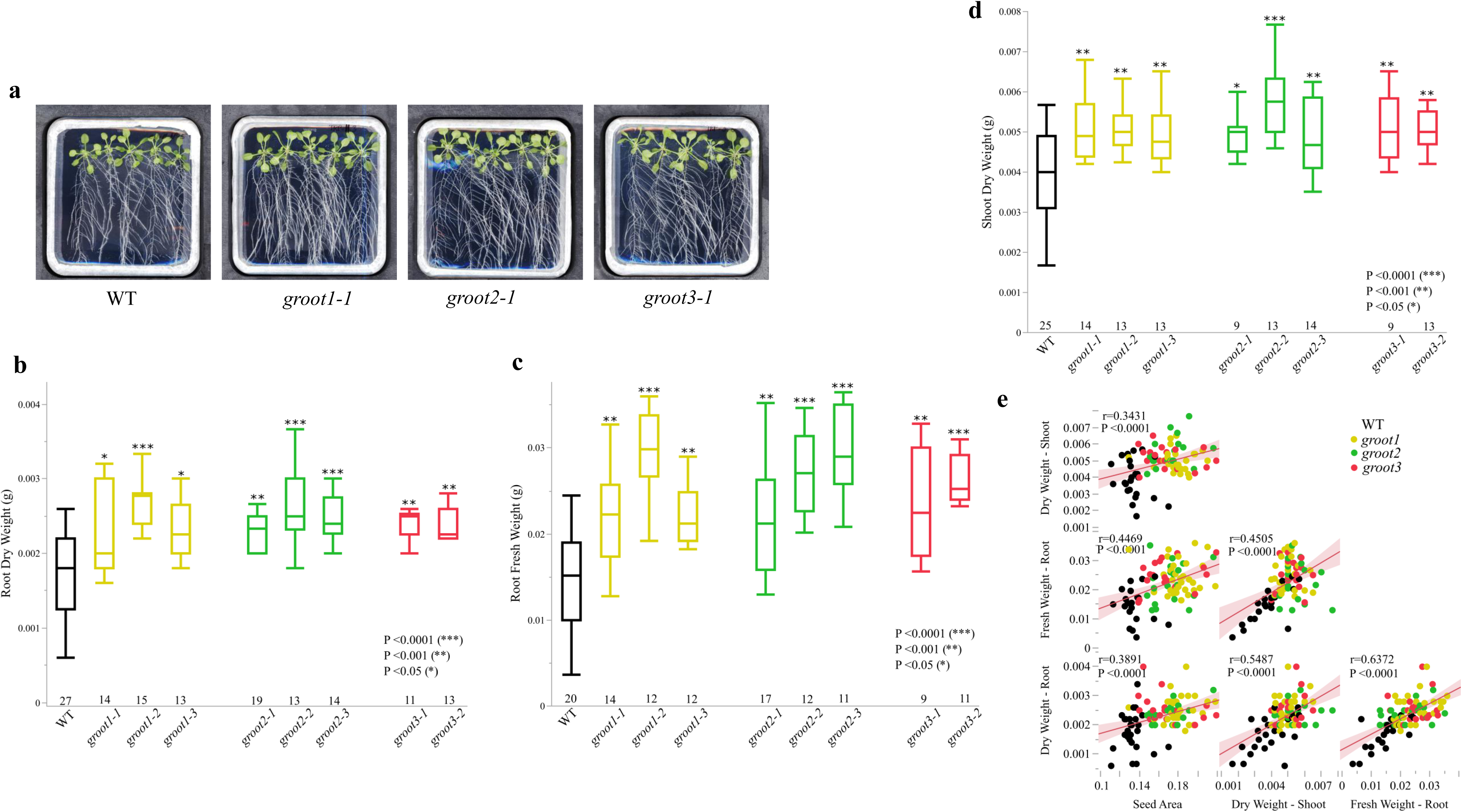
*groot* mutants show increased plant biomass and seed area. (a) Growth patterns of wild-type (WT) Col-0 and T-DNA lines for the three *GROOT* genes after 21 days on ½ MS plates. (b-d) Boxplot of root dry weight (b), root fresh weight (c), shoot dry weight (d) of Col-0 WT and T-DNA lines for *GROOT1*, *GROOT2*, and *GROOT3* at 21 days after planting. Color-coded by gene; Statistically significant (T-test) indicates by asterisks. (e) Correlations between WT and T-DNA lines for root dry weight, root fresh weight, shoot dry weight, and seed area. In box plots, the horizontal line indicates the median value, the lower and upper edges represent the 25th and 75th percentiles, and the whiskers extend to the minimum and maximum values. Dots represent individual lines; Colors: T-DNA lines; correlation coefficients (r) and P-values provided in figure.

We then set out to characterize these genes and their mutant lines further. *GROOT1* (*AT3G19440*) encodes a protein that belongs to the Pseudouridine synthase family. Members of these family have been shown to be enzymes catalyzing the conversion of pseudouridine (Ψ) to uridine (U) (Song et al. 2020). To confirm our initial results that we had obtained with one T-DNA mutant, we acquired three additional, independent T-DNA lines, in whose *GROOT1* was disrupted by T-DNA insertions in different parts of the gene including promoter, exons and introns (Supplementary File 8). We evaluated biomass changes compared to the wildtype (WT) reference line (Col-0) for root fresh weight and dry root weight after 21 days of growth under long-day conditions on MS plates (Fig. 3a). Like the first T-DNA line, all T-DNA lines exhibited significantly higher root fresh-weight compared to WT (P < 0.05). The mean percentage increase root in fresh weight was 61% compared to WT (Fig. 3b). Additionally, we also evaluated whether the increase in fresh weight went along with an increase in dry weight. The average increase in root dry weight biomass for these three T-DNA lines of *GROOT1* was 34%, with all lines displaying statistically significant differences at P < 0.05 compared to WT (Fig. 3b, c). All three T-DNA lines also displayed significantly higher shoot dry weight compared to the wild type (WT) (P < 0.05) (Fig. 3a, 3e). Overall, these three *GROOT1* T-DNA lines exhibited an average of 26% increase in shoot dry weight compared to the WT (Fig. 3d, Supplementary File 9). Overall, this confirmed that the *GROOT1* gene is involved in limiting plant growth.

*GROOT2* (AT3G19590) encodes a protein that belongs to the *BUB3* (*BUDDING UNINHIBITED BY BENZYMIDAZOL*) family. Members of this family have been shown to be involved in the spindle assembly checkpoint and gametophyte development (Lermontova et al. 2008). Again, we examined multiple T-DNA lines (Fig. 3a-b) that disrupted the *GROOT2* gene and assessed root fresh and dry weight biomass compared to the WT. Our findings revealed significant (P < 0.05) increases in biomass accumulation for root fresh weight, root dry weight biomass in all T-DNA lines compared to the WT (Fig. 3b, c). For root fresh weight, the T-DNA lines exhibited an average increase of 71% in mass compared to the WT (Fig. 3b). Similarly, root dry weight also showed a significant average increase of 37% compared to the WT for the T-DNA lines (Fig. 3c). There was also a significant elevation in shoot dry weight biomass in the *groot2* T-DNA lines compared to the WT, with an average increase of 30% (Fig. 3d, Supplementary File 9). Overall, this confirmed that the *GROOT2* gene is involved in limiting plant growth.

The *GROOT3* gene (AT3G19630) encodes a protein belonging to the *Radical SAM superfamily.* Members of this family have been shown to encode for enzymes that utilize SAM (S-adenosyl-L-methionine) to initiate radical reactions through liberation of the 5′-deoxyadenosyl (5′-dAdo) radical (Holliday et al. 2018; Hoffman et al. 2023). We identified two distinct T-DNA lines disrupting the function of this gene from the SALK T-DNA database (Fig. 3a-b), enabling us to investigate its impact on root fresh and dry weight biomass accumulation. Like the observations with *groot1* and *groot2* lines, both *groot3* T-DNA lines demonstrated significantly higher root fresh and dry weight biomass (Fig. 3a-d). On average, the *groot3* T-DNA lines exhibited a 66% increase in root fresh weight (Fig. 3b) and a 38% increase in root dry weight (Fig. 3c) compared to the WT. This underscores the significant role of *GROOT3* in limiting biomass accumulation in both root fresh and dry weight tissues. Like the other *groot* mutant lines, the *groot3* mutant lines also exhibited analogous patterns of significant shoot dry weight biomass increase compared to the wild type (Fig. 3a & 3d, Supplementary File 9), showing an average of 29% increase in shoot dry weight compared to WT. Overall, this confirmed that the *GROOT3* gene is involved in limiting plant growth.

As we had observed larger root and shoot mass in all *groot* mutant lines, we wanted to test whether *groot* seeds were larger as well. We therefore investigated the seed area (mm^2^) of WT and T-DNA lines for all three *GROOT* genes, aiming to comprehensively understand how mutant lines manifest and reveal the relationship between crucial life history traits and differences in biomass accumulation. We found a robust positive correlation between seed area and root dry weight (r = 0.39, P < 0.0001), root fresh weight (r = 0.45, P < 0.0001), and shoot dry weight (r = 0.34, P < 0.0001) (Fig. 3e). This result suggests that the increase in biomass accumulation goes along with increases in seed size among the *groot* mutant lines. Overall, these findings support that the three *GROOT* genes are not determining trade-offs between root and shoot growth or increases in seed size, but rather act as general growth limiters. This observation also hints at the potential evolutionary significance of altering multiple traits simultaneously, without experiencing negative trade-offs between different aspects of plant growth and development. It suggests an adaptive strategy where plants optimize resource allocation to maximize overall fitness, a concept central to evolutionary biology and ecological adaptation (Grime and Pierce 2012; Ackerly et al. 2000; Anderson et al. 2011; Smith 1978).

### The *GROOT* Genes are Associated with Different Growth-Related Processes

To further investigate the role of the *GROOT* genes, we investigated their expression pattern across different cell and tissue types using available published data. We first examined the expression patterns across different organs and developmental stages using the data from Klepikova et al. (2016). Our findings revealed that all three *GROOT* genes are expressed in various tissue types and developmental stages, though their expression levels varied. Notably, *GROOT3* exhibited higher expression levels across all tissue types and developmental stages compared to *GROOT1* and *GROOT 2* (Fig. 4a-c). While all three genes were expressed in common developmental stages and tissue types such as the root, seed, shoot apex, and shoot system, each gene displayed unique expression characteristics. To further analyze their expression patterns in the root at a single-cell resolution, we utilized published single-cell root data from Shahan et al. (2022). This analysis revealed additional variations in specific cell type expression patterns (Fig. 4d). For instance, *GROOT1* exhibited higher expression levels in the xylem pole pericycle, phloem pole pericycle, and the quiescent center. In contrast, *GROOT2* displayed an elevated expression in the lateral root cap, while *GROOT3* showed a higher expression in the quiescent center (Fig. 4d). These genes are also expressed in both dividing (meristem) and maturing tissues in the root. However, while all three genes are expressed in the dividing tissues, their expression in maturing tissues varies. For instance, *GROOT1* is expressed in the xylem pole pericycle maturing tissue, *GROOT3* is expressed in all maturing tissues, and *GROOT2* is restricted to the lateral root cap maturing tissue (Fig. 4e). These observations suggest that these three genes might play unique roles at both tissue and cellular levels, indicating diverse functions and regulatory mechanisms.

**Figure 4:**
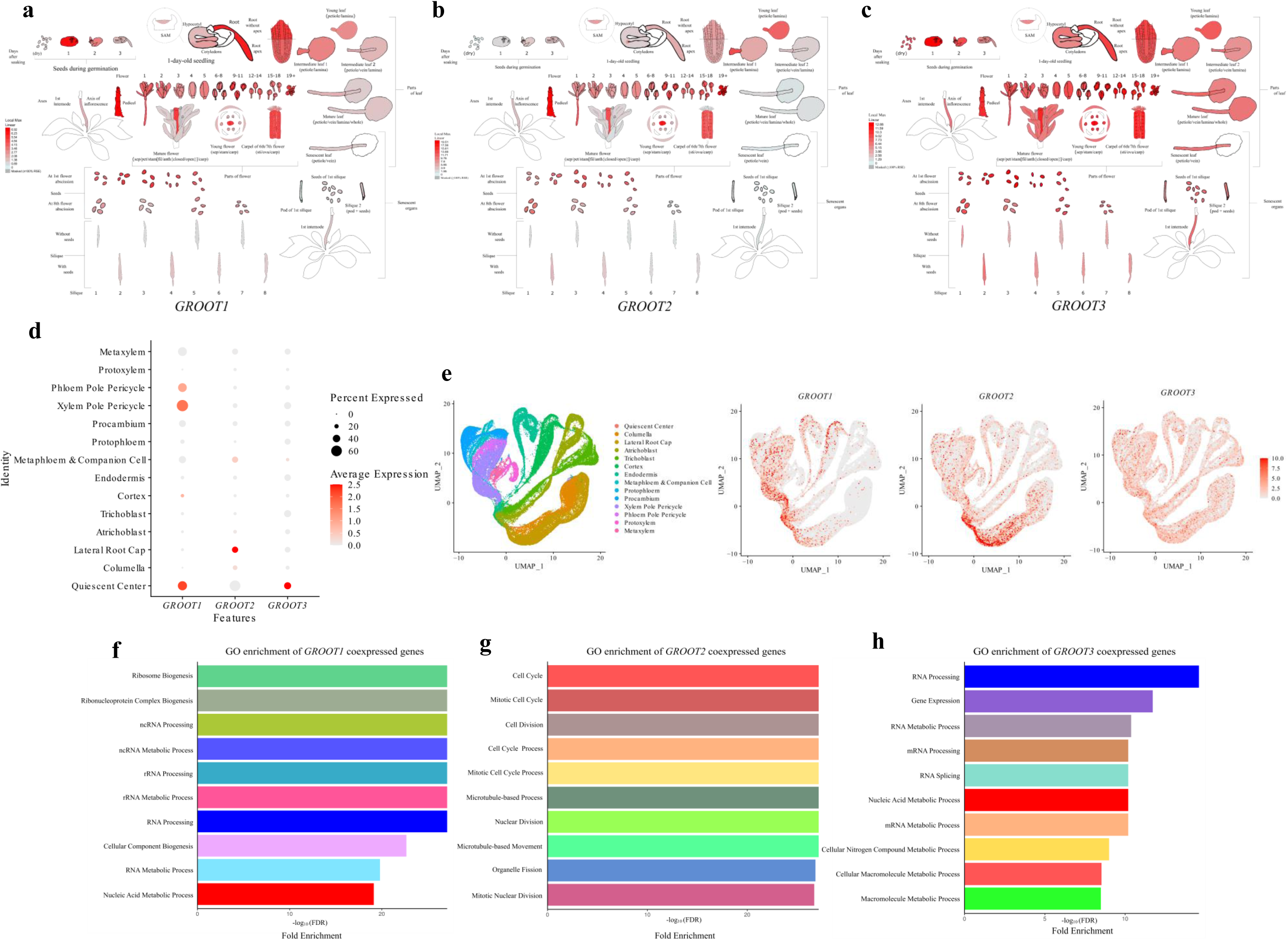
*GROOT* gene expression patterns and coregulated genes suggest diverse biological functions. (a-c) Tissue expression of *GROOT1*, *GROOT2*, and *GROOT3* across different organs and developmental stages according to Klepikova et al. (2016). (d,e) Single cell RNAseq root expression data according to Shahan et al. (2022 (d) y-axis: cell types; x-axis: genes; dot color indicates intensity of average expression level and dot size indicates percentage of cells within cell type expressing the gene (e) UMAP plots for *GROOT* genes. (f-h) GO enrichment for the 200 co-expressed genes of *GROOT* genes according to AttedII database (https://atted.jp/). Y-axis: GO category for biological processes; x-axis: Represents the fold enrichment, with the bar length indicating the significance of enrichment as measured by - log10(FDR) values.

To obtain a glimpse in their biological function, we then explored co-expressed genes of *GROOT1*, *GROOT2*, and *GROOT3* by examining GO enrichment of co-expressed genes. This analysis aimed to uncover the specific biological functions and processes these co-expressed genes are involved in at cellular and tissue-specific levels. By identifying enriched GO terms, we sought to understand the roles and interactions of these genes within the broader biological context, revealing how they contribute to essential processes like cell division, RNA processing, and ribosome biogenesis. This approach also allowed us to generate hypotheses about the regulatory networks and pathways these genes participate in, providing deeper insights into their contributions to growth, development, and overall biomass production. Overall, the GO enrichment analysis revealed distinct functional categories, which might be connected to their association with increased biomass production (Fig. 4f-h). Consistent with its annotation as Pseudouridine synthase family member, *GROOT1* co-expressed genes are enriched for GO categories such as ribosome biogenesis, RNA processing, and RNA metabolic processes, highlighting its likely role in ribosome production and RNA metabolism, which are crucial for protein synthesis and overall cellular growth (Fig. 4f) and have been shown to be involved in natural variation of root growth rate (Slovak et al. 2020). Consistent with its annotation as a cell-cycle related gene, *GROOT2* co-expressed genes are enriched in GO categories associated with the cell cycle, cell division, and nuclear division, indicating its involvement in regulating cell proliferation and ensuring accurate chromosome segregation during mitosis, processes vital for sustaining rapid growth and biomass accumulation (Fig. 4g). Consistent with some members of the Radical SAM superfamily having a role in RNA processing*, GROOT3* co-expressed genes are enriched in GO categories related to RNA processing, gene expression, and RNA splicing, suggesting its role in gene expression and RNA processing, essential for the efficient functioning and growth of cells (Fig. 4h). These functional roles might suggest that *GROOT3* contributes to the regulation of gene expression, *GROOT2* to cell division and growth, and *GROOT1* to protein synthesis, all of which are essential processes for biomass production. Therefore, the high expression and functional roles of these genes likely underpin their association with increased biomass, as they collectively enhance the cellular and molecular mechanisms that drive growth and development.

### The linkage disequilibrium pattern of *GROOT* alleles suggests a potential for coinheritance

The attribution of all three *GROOT* genes that are involved in limiting biomass accumulation to a single GWAS peak intrigued us. We therefore wanted to elucidate whether natural alleles of these genes are connected to one another. For this, we conducted an analysis of Linkage Disequilibrium (LD) of all three *GROOT* genes with the top GWAS SNP (6772287) in this region within a 70kb window. Overall, the LD between the GWAS SNP and other SNPs in this region was relatively low, with less than 1.1% of SNPs exhibiting an LD r^2^ value over 0.2, suggesting a weak association of LD with the top GWAS SNP within the region. However, within the *GROOT* genes we identified several SNPs displaying strong LD with the top GWAS SNP. Notably, a SNP (6810046) in the *GROOT2* gene exhibited the highest LD (r^2^ = 0.66) with the top GWAS SNP (6772287). Similarly, a SNP (6745394) in *GROOT1* displayed a high LD (r^2^ value) of 0.62 with the top GWAS SNP. The SNP in *GROOT3* displaying the highest LD with the top GWAS SNP was SNP 6815838 showing an LD of 0.48.

This LD pattern prompted us to investigate the relation of *GROOT* variants (as defined by their SNPs in LD with the top GWAS hit) with the root biomass trait. To achieve this, we examined the pattern of sequence polymorphism for these gene positions within the natural accessions used in our study (https://tools.1001genomes.org/polymorph/). We grouped all natural accessions based on whether they had reference alleles or non-reference alleles at each position and then analyzed the relationship between these SNP patterns and the biomass trait. Our analysis revealed that accessions with reference alleles in *GROOT1* were linked to higher biomass values (Fig. 5a). For both *GROOT3* and *GROOT2*, accessions with non-reference alleles exhibited significantly higher biomass values compared to those with reference alleles (Figs. 5b-c). Given the distinct and potentially complementary molecular roles of the three *GROOT* genes, we then wanted to ask whether a combination of biomass increasing alleles would be associated with a non-additive increase compared to single or double allele combination. For this, we grouped accessions according to their *GROOT* allele combination. Consistent with the idea that these alleles might act in a synergistic fashion, we found that accessions with a combination of all three biomass increasing alleles in average showed *55.6%* increase of TRP compared to accessions that didn’t contain any biomass increasing *GROOT* alleles (Fig. 5d). In fact, accessions that only contained one or two biomass-increasing GROOT alleles, didn’t display significantly higher biomass than accessions containing none of the three biomass-increase associated GROOT alleles (Fig. 5d). This might indicate a synergistic effect of all three *GROOT* alleles to lead to higher root biomass.

**Figure 5:**
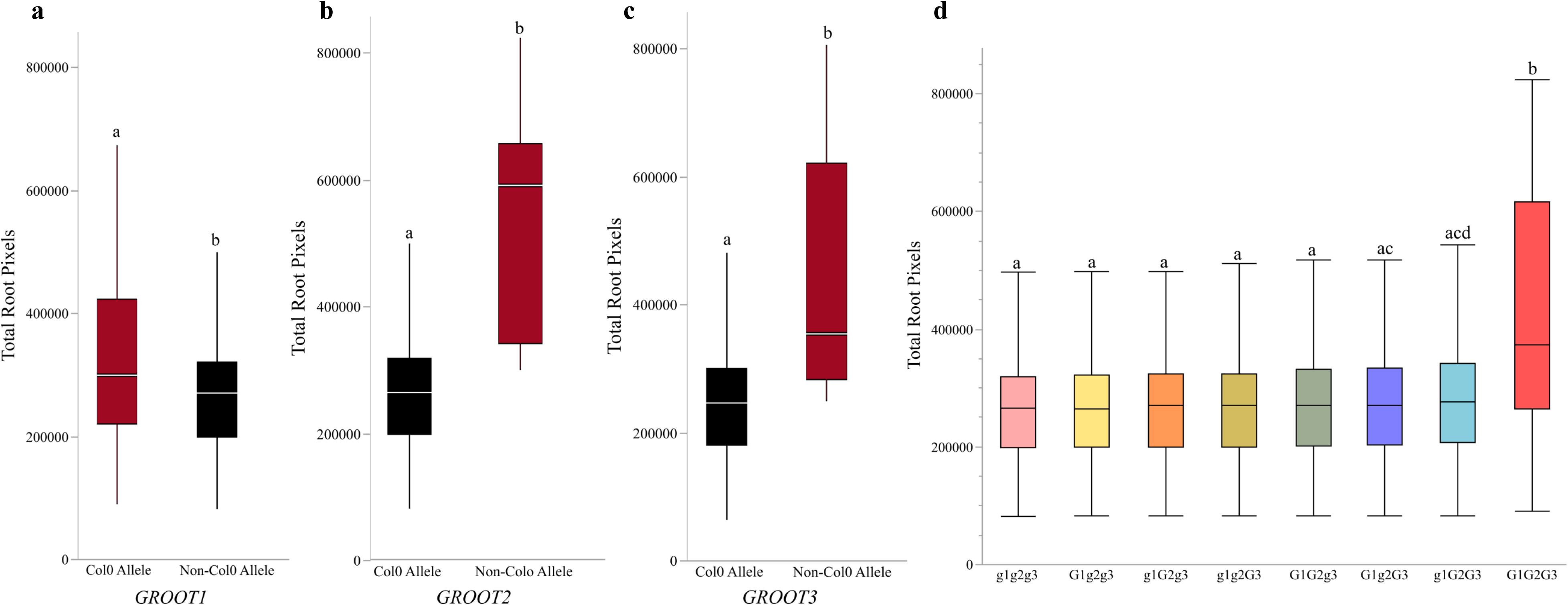
*GROOT* gene SNPs display a robust association with biomass accumulation. (a-c) Boxplot TRP associated with *GROOT* gene variants and their association with. Y-axis represents TRP value; X-axis represents types of variants (Col-0 allele and non-Col-0 allele); SNP position (a) 6745394 for *GROOT1*, (b) 6810046 for *GROOT2,* (c) 6815838 for *GROOT3*. Significant differences (P < 0.05) were tested using a T-test and are indicated by letters above the error bars. (d) TRP in accessions grouped by major and minor alleles of *GROOT1*, *GROOT2*, and *GROOT3*. X-axis: unique allele combination; y-axis: TRP of accessions in this group. Letters denote significance levels.

Overall, our analysis demonstrated that specific SNPs within the *GROOT* genes exhibit strong linkage disequilibrium (LD) with the GWAS top SNP, suggesting that these loci may have been targets of positive selection. This high LD implies that allelic variants at these SNPs are co-inherited more frequently than would be expected by chance, indicating a selective sweep or historical selection pressure that has preserved these advantageous genetic combinations. Moreover, the SNPs in these genes showing high LD were also associated with high biomass production, with most of the high biomass-producing accessions overlapping. The significant LD observed suggests that these alleles contribute to the adaptive variation in biomass accumulation, reflecting a potential role in the evolutionary fitness of the populations. This interconnectedness suggests the importance of these SNPs in shaping the genetic architecture underlying biomass production, offering insights into the evolutionary dynamics driving trait variation in these genes. These findings underscore the potential of these loci as key targets for understanding the genetic basis of biomass yield and for leveraging advantageous genetic variants in related research.

### Local climate dynamics drive interactions between root growth patterns and genetic variants, shaping plant adaptation strategies

Root growth rate increases with temperature (Gaillochet et al. 2020; Lee et al. 2023) and we therefore wanted to further explore the link between the allelic variants of the confirmed GWAS hits and local climate conditions. We therefore investigated the relation of local conditions, such as temperature or precipitation, and the allelic variant (SNP Chr3 at 6772287) that was associated with all three of our biomasses linked *GROOT* genes. We first examined the relationship between bioclimatic variables, and all studied natural accessions grouped by their allelic variants for the GWAS SNP (6772287). We identified several environmental factors significantly associated with non-reference allele variants. Accessions with non-reference alleles exhibited higher values for factors related to temperature, including mean diurnal temperature (P = 0.001), isothermality (P = 0.0002), and mean temperature of the driest quarter (P = 0.0030, Supplementary Figure 2). This might indicate a significant association between high temperatures and enhanced root biomass. The interpretation of these findings indicates that specific allelic variants may confer an adaptive advantage under certain climatic conditions, particularly higher temperatures, potentially leading to increased root biomass. This adaptation could be crucial for plant survival and productivity in varying environmental conditions, highlighting the importance of these genetic variants in climate resilience.

To further investigate this hypothesis, we selected extreme natural accessions from the TRP distribution, marked in red (Fig. 6a). When selecting accessions for this study, we ensured representation from both reference allele and non-reference allele groups. This approach allowed us to assess the performance of reference allele-associated (Col-0) accessions in high-temperature conditions relative to accessions containing the non-reference (non-Col-0) allele (Supplementary File 10). After growing the accessions to both elevated (28°C) and regular lab-grown temperatures (22°C) for 21 days, we analyzed the fresh and dry weights of root and shoot tissue, observing variable growth patterns at elevated temperatures. For instance, BÃ¥5-1 containing reference Col-0 allele and IP-Pie-0 containing non-Col-0 allele displayed distinct growth patterns under elevated temperatures (Fig. 6b). We then analyzed the effects of genotype (G), temperature (E), and their interaction (GxE) on the traits studied. Within this framework, significant G signifies divergence in root traits between reference allele and non-reference allele lines, significant E indicates temperature-driven plasticity, and significant GxE suggests variation in plastic responses between lines containing different alleles. While the increased biomass of root and shoot growth in non-Col-0 allele-containing accessions compared to Col-0 allele-containing accessions persisted, the rate of increase also significantly differed between the two groups of accessions. Specifically, the non-Col-0 allele containing accessions exhibited a more pronounced growth response compared to the Col-0 allele containing accessions (Fig. 6, Supplementary File 10. Consequently, significant effects of genotype (G, P < 0.0001) and temperature (E, P < 0.0001) were observed for both root fresh weight and dry weight. This indicates that both the genetic makeup of the plants and the environmental conditions (in this case, temperature) play crucial roles in determining the biomass of the roots. Interestingly, the genotype-by-environment interaction (GxE) was non-significant for root fresh weight (P = 0.6766) but significant for root dry weight (P = 0.0132, Fig. 6c-d). This suggests that while the immediate water content (fresh weight) of the roots was not significantly influenced by the interaction of genotype and environment, the overall root biomass (dry weight), which is more direct measure of growth, was indeed affected by how different genotypes with varying allelic variants responded to different temperatures. Similar trends were noted for shoot fresh and dry weights. Both genotype (G) and temperature (E) had significant effects (P < 0.0001) on shoot biomass. Moreover, their interaction effects were also significant for both shoot fresh weight (P = 0.0091) and shoot dry weight (P = 0.0420) (Fig. 6e-f). These findings highlight the intricate nature of genotype-by-environment (GxE) interactions and their vital role in shaping plant growth responses to varying temperatures.

**Figure 6:**
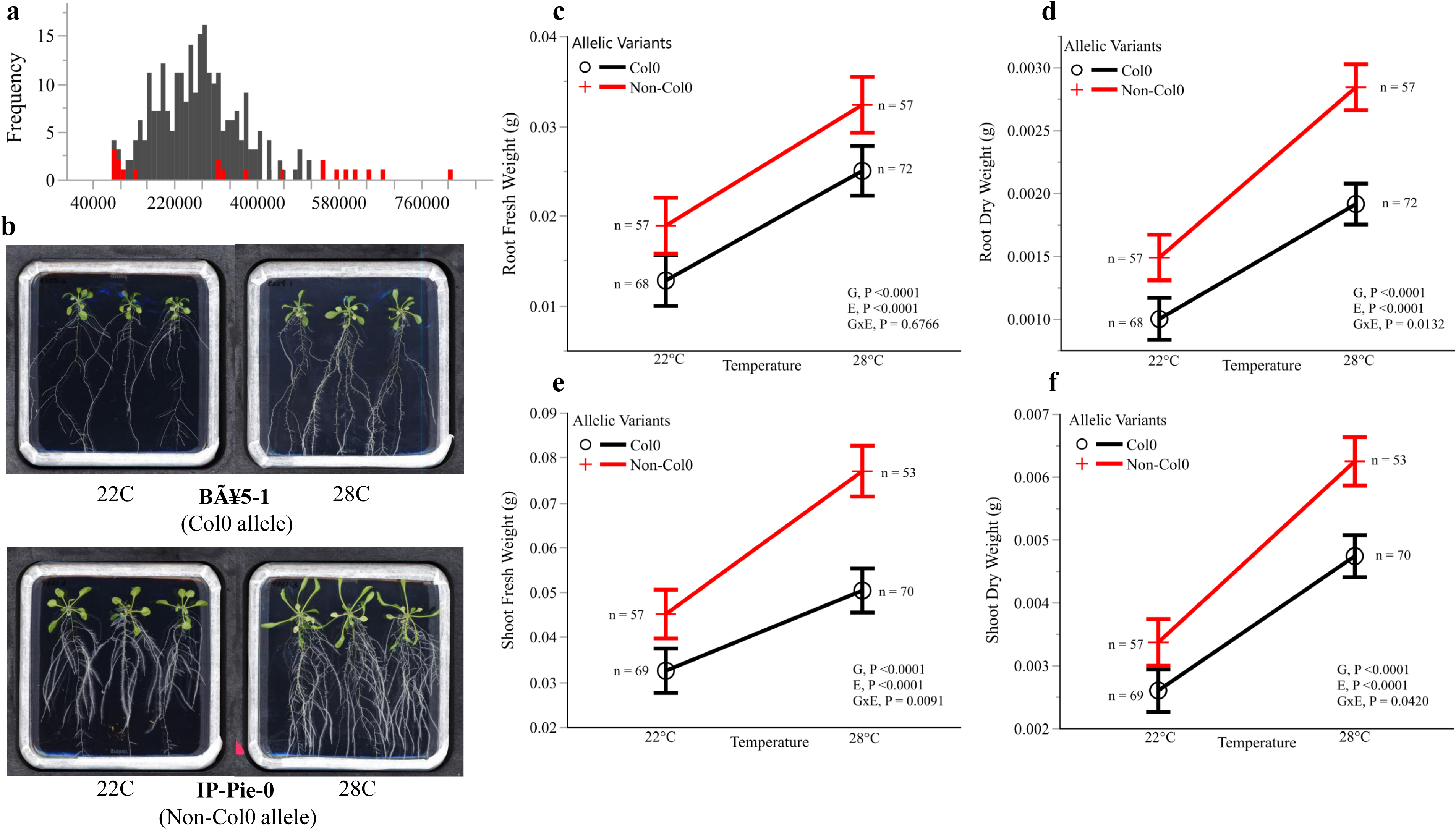
Root mass accumulation under elevated temperatures in natural accessions. (a) Position of accessions used for elevated temperature experiment (red) in TRP data distribution. Y-axis: frequency; X-axis: TRP. (b) Growth differences between two accessions with different SNP types at Chromosome three position 6772287 at 22°C and 28°C. Bå5-1 contains Col-0 allele; IP-Pie-0 contains non-Col-0 allele. Both lines show higher root mass at elevated temperatures. (c-f) Phenotypes of Col-0 and non-Col-0 allele accessions at 22°C and 28°C. (c) Root fresh weight, (d) root dry weight, (e) shoot fresh weight, (f) shoot dry weight. Significance levels for interaction terms are denoted in the figure (G: Genotype; E: Environment). Significance was determined using a two-way ANOVA followed by Tukey’s HSD test (p < 0.05). This experiment involved 22 accessions, with data collected from root and shoot tissues at 21 days after planting (DAP). These accessions were grouped into 12 with the Col-0 allele and 10 with the Non-Col-0 allele. The figure displays the total number of data points collected from each group.

Finally, we wanted to explore the effect of elevated temperature on the impact of the three *GROOT* genes. Like for the accessions, we gathered data from mutant lines and WT plants grown at both 22°C and 28°C. On day 7, primary root length (cm) showed substantial main effects for genotype (P < 0.0001), temperature (P < 0.0001), and a genotype-by-temperature interaction (P = 0.0031; Fig. 7a & 7c, Supplementary File 11). By day 14, we also noted accelerated primary root growth at the higher temperature (28°C) compared to the lower temperature (22°C) (Fig. 7b). Within the genotype-by-temperature interaction framework (GxE), genotype (G, P < 0.0001), temperature (E, P < 0.0001), and their interaction (GxE, P < 0.0001) significantly influenced primary root growth at day 14 (Fig. 7b & 7d, Supplementary File 11). Furthermore, shoot and root dry mass at day 21 under elevated temperature conditions revealed significant effects of genotype (G, P < 0.0001), temperature (E, P < 0.0001), and genotype-by-temperature interaction (GxE, P < 0.0001) on both root and shoot biomass (Fig. 7e-g, Supplementary File 11). It is worth noting that we observed reduced biomass accumulation at elevated (28°C) temperatures compared to regular (22°C) conditions. Specifically, under elevated temperature conditions, all T-DNA lines, including the wild type (WT), began flowering from day 17 on the plate. This shift towards reproductive growth post-flowering may significantly limit biomass accumulation post day 17. These observations might suggest that the three identified genes play a role in growth limitation under varying temperature regimes. This highlights the importance of genotype-by-environment (GxE) interactions in understanding how these genes regulate root growth, particularly in response to elevated temperatures.

**Figure 7:**
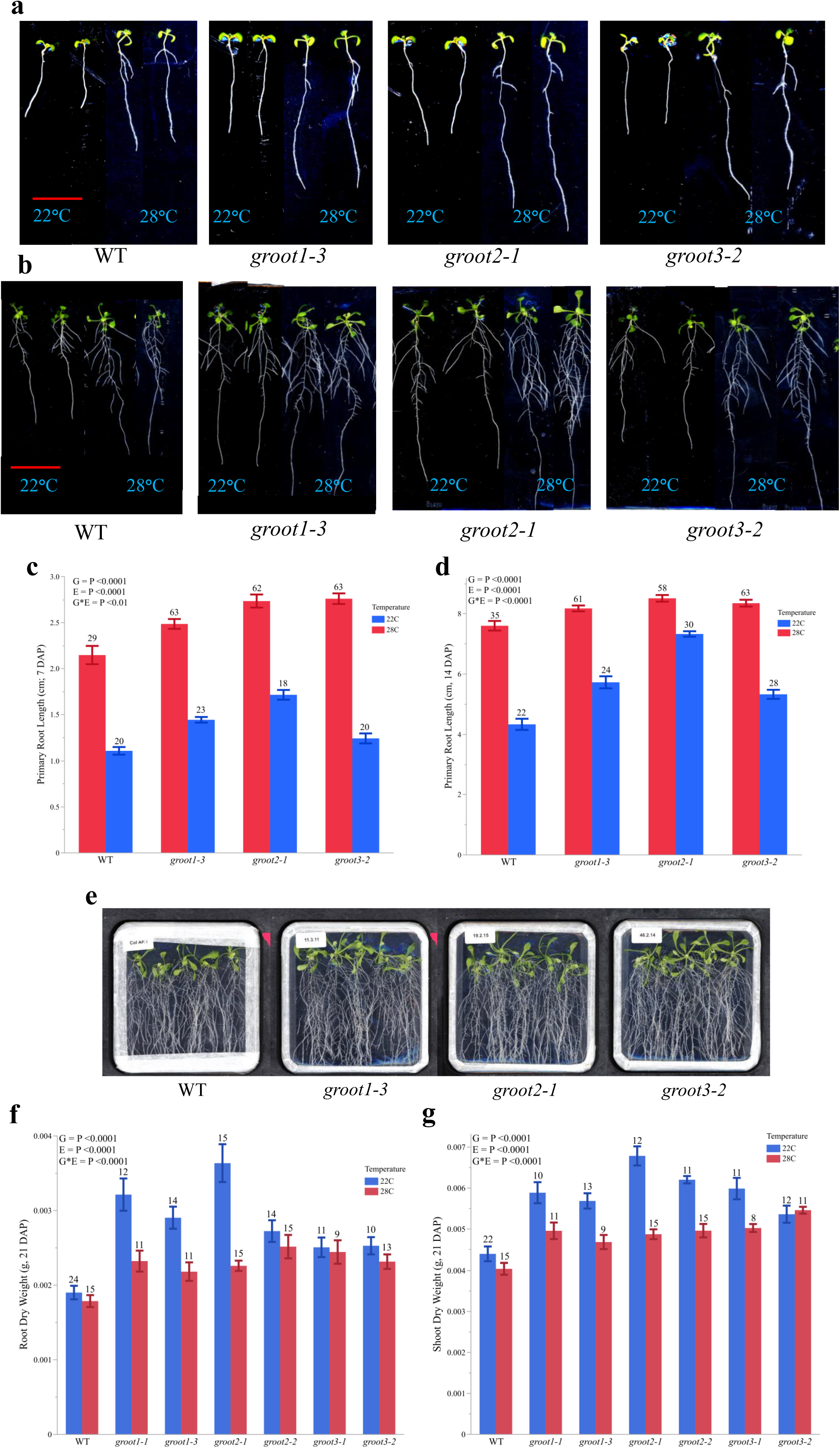
Plant biomass in *groot* mutants displays a significant genotype-by-environment (GxE) interactions. (a,b) Root growth patterns for WT and *groot1*, *groot2*, and *groot3* at 7 (a) and 14 (b) days after planting. (c,d) Root growth of *groot* mutants and WT at 7 (c) and 14 (d) days after planting. Y-axis: Primary root length (cm). Significance levels for interactions terms in figure (G: Genotype; E: Environment) impact root length. (e) WT and *groot* mutant plants at 21 days after planting at 28°C. (f-g) Shoot and root dry weight biomass response after 21 DAP to elevated temperature. Y-axis: root (f) and shoot (g) dry weight biomass. Significant GxE interactions drive biomass accumulation. Significance levels for interaction terms are denoted in the figure (G: Genotype; E: Environment). Significance was determined using a two-way ANOVA followed by Tukey’s HSD test (p < 0.05). The number of replicates for each T-DNA line and wild-type is indicated at the top of each bar.

## Discussion

In this study, we developed a high-throughput, deep learning-based technique called the Total Root Pixel (TRP) method for accurately estimating root biomass, particularly focusing on fine roots, which are often challenging for existing tools. This method simplifies the traditionally labor-intensive process by providing a non-destructive, image-based alternative, with a strong correlation observed between TRP estimates and actual biomass measurements (Fig. 1d). Using the UNet^++^ architecture, the TRP method effectively segments and quantifies root pixels from images of seedlings grown on Murashige and Skoog (MS) plates over 21 days. When applied to a GWAS, the TRP method identified several candidate loci, underscoring the polygenic nature of complex traits such as root biomass, in line with findings from other studies on plant development related traits (Weigel and Nordborg 2005; Rockman 2012; Courtois et al. 2013; Zurek et al. 2015). This tool represents a significant advancement in root phenotyping, offering a scalable solution for accelerating genetic research and breeding programs aimed at improving crop resilience. The development of such methods is crucial, especially as we face increasing environmental challenges, emphasizing the importance of tools that enhance our understanding of root system architecture and its genetic basis (Minervini et al. 2017; Smith et al. 2022).

Our study leveraged the TRP-based root biomass trait in a genome-wide association study (GWAS) to investigate the genetic architecture of root biomass, uncovering insights that align with previous research identifying multiple loci associated with complex traits such as root biomass across various plant species. For example, in *Arabidopsis thaliana*, comprehensive GWAS analyses have pinpointed several loci significantly influencing several life history traits (Atwell et al. 2010; Ristova and Busch 2014). Our findings also showed that the top five significant GWAS loci were predominantly associated with accessions from a specific geographical region. These accessions, carrying mutations at the highest GWAS peaks, consistently exhibited higher root biomass (Fig. 2a-c), suggesting that selection pressures in this region may favor these loci, potentially as an adaptation to local environmental conditions. This pattern of regional adaptation is well-documented in the literature, where traits such as drought tolerance and flowering time have been shown to be influenced by local environmental factors, leading to region-specific selection (Fournier-Level et al. 2011; Lasky et al. 2012; Savolainen et al. 2013). For example, environmental factors like soil composition, moisture availability, and nutrient levels may drive the selection of genotypes with enhanced root biomass accumulation. These observations align with the understanding that complex traits like root biomass are polygenic, with multiple loci contributing small but cumulative effects, as highlighted in studies on root development and other complex traits (Weigel and Nordborg 2005; Rockman 2012; Courtois et al. 2013). Thus, our study adds to the growing body of evidence emphasizing the need to consider both genetic and environmental factors when investigating complex traits. Recognizing the polygenic nature and regional specificity of these traits is essential for devising strategies to improve crop resilience and optimize growth across diverse environments.

In this study, we identified and characterized three closely linked genes—*GROOT1*, *GROOT2*, and *GROOT3*—that significantly regulate plant biomass accumulation (Fig. 3). These genes, located in close proximity on chromosome 3, exhibit strong linkage disequilibrium with SNPs identified in our GWAS, indicating a coordinated influence on root and shoot biomass traits. Disruption of these genes in multiple T-DNA mutant lines resulted in substantial increases in both root and shoot biomass, with no observable trade-offs in other critical life history traits (Fig. 3). The absence of trade-offs, along with additive effects across multiple traits, suggests that *GROOT1*, *GROOT2*, and *GROOT3* function as general growth limiters. This finding indicates that these genes are not simply reallocating resources between different growth processes but instead broadly restricting growth potential across the plant. Such a mechanism could provide a selective advantage by enabling plants to optimize resource allocation and maximize overall fitness in environments where competition for resources is intense (Grime and Pierce 2012; Smith 1978). Additionally, the close physical proximity of these genes, coupled with high linkage disequilibrium, suggests that they may operate as a coordinated cluster, contributing to a unified regulatory mechanism that limits growth across multiple traits. This clustering and strong linkage disequilibrium might reflect an evolutionary strategy, where these genes have been selected together to ensure robust control over developmental processes, allowing plants to thrive under diverse environmental conditions (Weigel and Nordborg 2005; Anderson et al. 2011). Overall, our findings provide new insights into the genetic regulation of plant growth and suggest that these genes play a critical role in optimizing resource allocation and overall plant fitness, particularly in response to varying environmental conditions.

Single nucleotide polymorphisms (SNPs) identified in our GWAS are associated with a significant response to elevated temperatures. Specifically, accessions carrying the SNP at position 6772287 on chromosome 3 showed markedly higher shoot and root biomass under elevated temperatures (28°C), both in fresh and dry weight, compared to those grown under standard conditions (22°C; Fig. 6). This might suggest that accessions with this SNP might be regularly exposed to temperature fluctuations in their native environments, leading to the selection of this allele for improved adaptation to higher temperatures. The relationship between SNPs and temperature adaptation is an area of growing interest. Previous studies have shown that certain SNPs can be linked to improved thermal tolerance in plants. For example, Hancock et al. (2011) demonstrated that specific SNPs were associated with heat tolerance in *Arabidopsis thaliana*, where plants with these alleles exhibited better growth and survival under heat stress. Similarly, in rice, studies have identified SNPs associated with enhanced heat tolerance, where alleles at these loci have been shown to contribute to improved grain yield under high-temperature conditions (Ye et al. 2012). These findings underscore the vital role of specific genetic variants in enabling plants to endure and prosper in elevated temperatures. Our research extends this understanding by suggesting that the SNP at position 6772287 on chromosome 3 may be instrumental in facilitating temperature adaptation, making it a promising target for future research focused on enhancing crop resilience in the face of climate change. In line with that, *groot* mutant lines revealed a similar adaptive response, with increased primary root length observed under elevated temperatures on days 7 and 14. However, by day 21, regular temperature conditions resulted in greater dry mass accumulation, likely due to early flowering triggered by elevated temperatures, as described by Blázquez et al. (2003) and Balasubramanian et al. (2006). This early flowering may have limited vegetative growth, contrasting with the sustained growth seen in SNP-containing accessions. These observations might suggest that the *GROOT* genes may play a role in temperature adaptation, potentially playing a role for root development in warmer climates.

In summary, our findings suggest that the SNP at chromosome 3 position 6772287 may be an important genetic marker for temperature adaptation in plants. The differential growth responses observed between SNP-containing accessions and *groot* mutant lines under elevated temperatures further underscore the complexity of plant responses to different environments and the potential for specific genetic loci to confer adaptive advantages. In addition, our study highlights genetic strategies to increase plant biomass by mutating the orthologues of *GROOT* genes in crop species. In contrast to previous genetic interventions that required transgenes, *groot* knockout are compatible with modern gene editing strategies. Therefore, future studies exploring the molecular mechanisms underlying our observations and investigating the broader applicability of these findings in other plant species and environmental contexts promise to provide insights into plant adaptation as well as crop improvement strategies.

## Materials and methods

### Plant materials and growth conditions

Seeds of *Arabidopsis thaliana* were surface sterilized by placing them in open 1.5 ml Eppendorf tubes within a sealed environment with chorine gas. Chlorine gas was generated from a solution comprising 10% sodium hypochlorite (130 ml) and 37% hydrochloric acid (3.5 ml) that was kept in the sealed container with the seeds for an hour. Afterward, for stratification, the seeds underwent water imbibition followed by a 4-day period at 4 °C in darkness. After that the seeds were sown directly on ½ MS agar plates with a pH of 5.7, comprising of 1% (w/v) sucrose and 1% (w/v) phytagel sourced from Sigma (www.sigmaaldrich.com/), plated within 12 cm × 12 cm square plates from Greiner (https://shop.gbo.com/). The plants were grown under long-day conditions (16 hours light/8 hours dark) in a walk-in growth chamber maintained at 21°C, with a light intensity of 50 μM and 60% humidity. Nighttime temperatures were reduced to 16°C. Seedlings were allowed to grow for 21 days before imaging, with three seedlings per plate for each line, and each line was replicated four times (plates served as the unit of replication).

### High-throughput image-based deep learning (UNet^++^) pipeline for root biomass phenotyping

After allowing the seedlings on the plates to grow for 21 days, we captured images of the plates using CCD flatbed scanners (EPSON Perfection V600 Photo, Seiko Epson CO., Nagano, Japan). These images were used to quantify total root pixel counts using a bespoke deep learning-based approach termed the Total Root Pixel (TRP) method. The method operates through several stages, encompassing image pre-processing, UNet^++^ model training, prediction generation, post-processing, and, ultimately, phenotyping. Comprehensive explanations of each step are provided in the following sections, while an overview of the entire procedure is depicted in Figure 1b. Each plate image, containing three Arabidopsis seedlings, was manually cropped to separate individual seedlings. Overlapping roots from different seedlings were carefully delineated and masked to prevent analysis errors. The training dataset was prepared by manually annotating roots and the surrounding background in the images using the *LabelMe* tool (Russell et al. 2008). To ensure comprehensive detail capture without losing boundary information, high-resolution images and their annotations were split into 512x512 pixel patches with a 16-pixel overlap. The training set, comprising 40 images with cropped seedlings, was randomly selected from the plate images.

We then utilized the UNet^++^ architecture for root segmentation enhanced by a ResNet18 encoder, known for its efficiency in handling detailed images (Zhou et al. 2018; Ronneberger et al. 2015). The model used a *sigmoid* activation function, which is particularly effective for binary segmentation tasks. Training was conducted over 40 epochs with an initial learning rate of 0.0001, utilizing the *Adam* optimizer. The model’s efficacy was evaluated using the Intersection over Union (IoU) metric, ensuring that only the best-performing model was retained for future predictions. Additionally, data augmentation strategies were integrated to bolster the model’s robustness and adaptability across varied imaging conditions (Shorten and Khoshgoftaar 2019).

In the prediction phase, individual seedling images were cropped, split into patches, and segmented using the trained UNet++ model. Segmented patches were then stitched to reconstruct the original layout of the cropped seedlings. A custom Python script utilizing the OpenCV library was developed to refine segmentation by applying thresholding techniques and identifying connected components. The largest component was targeted as the root, with noise reduction performed by excluding small irrelevant components, ensuring high fidelity in the analyses. The GitHub repository at https://github.com/idelly007/TRP-Total-Root-Pixel-Pipeline contains the scripts developed for the TRP pipelines.

Employing the developed method, the total root pixels (TRP) were quantified for 12 seedlings per accession. Subsequently, to validate the accuracy of the biomass estimate derived from TRP image pixel calculations pipeline, we grew a representative subset of the accessions employed in the estimation process to acquire data on root fresh weight. To achieve this, a total of 72 accessions were randomly selected and cultivated on MS plates for a duration of 21 days. After this growth period, the roots and shoots were carefully separated into distinct collection tubes. After ensuring the removal of any excess moisture from the growth media by employing Kim wipes, the separated plant roots were promptly weighed. The assessment of fresh weight biomass for the root tissue was conducted using a Toledo scale. Then we checked the correlation coefficients (r) between the estimates of root biomass done by the developed TRP method and the actual harvest of the root tissue.

### Broad sense heritability (H^2^) calculation

The broad-sense heritability (H^2^ = V_G_/V_P_) was calculated using the TRP data from 264 accessions. We used h2boot software fitting one-way ANOVA among individual lines with 1000 bootstrap runs (Phillips and Arnold 1999). Broad-sense heritability represents the proportion of phenotypic variation (V_P_) attributed to genetic variation (V_G_), estimated from the between-line phenotypic variance.

### Genome-Wide Association (GWA) mapping with the TRP data

We performed a Genome-Wide Association Study (GWAS) using the mean TRP values of 264 accessions. We conducted Genome-Wide Association (GWA) mapping using fully imputed SNP data from the 1001 Arabidopsis database (https://1001genomes.org/) with the Efficient Mixed Model Analysis (EMMA) mixed model algorithm as outlined by (Kang et al. 2010), integrated within the PyGWAS software framework. SNPs with minor allele counts (MAC) of 10 or more were considered. The significance of SNP associations was evaluated at a 5% False Discovery Rate (FDR) threshold (P <0.05) computed using the Benjamini-Hochberg method to address multiple testing (Benjamini and Yekutieli 2001).

### Analysis of Linkage disequilibrium (LD)

To assess Linkage Disequilibrium (LD) (r^2^) at the GWAS peak, we employed plink 1.9 (Purcell et al. 2007) with a window size of 70 kb (“–ld-window-kb 70”). The significance of r^2^ was determined using the 95th percentile (P <0.05) across the window.

### T-DNA insertion lines and phenotyping

Based on the GWAS results, we acquired T-DNA insertion lines for 51 candidate genes from ABRC (https://abrc.osu.edu/). The list of these genes and their associated details are in Supplementary File 1. Upon receipt of the lines, our first step was to genotype them to validate their homozygous insertion status. The PCR genotyping primers for each T-DNA line are collected using the Salk T-DNA primer design database (http://signal.salk.edu/tdnaprimers.2.html). Subsequently, we conducted phenotypic assessments on the homozygous T-DNA lines for root biomass and primary root length by growing them on ½ MS plates. To test the phenotype of the T-DNA lines, as a WT control, we initially utilized wild-type (WT) (Col-0) samples obtained from three distinct sources.

We grew the T-DNA lines and wild-type (WT) plants for 21 days under long-day conditions before harvesting them. The seeds were directly sown onto MS plates and stratified in darkness at 4°C. Upon transferring them to light, we initiated the planting day count. Each plate contained 5 seeds, and to minimize potential biases, we rotated the plates within the growth chamber every 3 days. Plate scanning occurred on days 7, 14, and 21. Primary root length was measured using ImageJ software based on images captured on days 7 and 14. On day 21, seedlings from each plate were harvested and pooled into pre-weighed Eppendorf tubes to ensure accurate fresh and dry weights. Due to the small size and weight of Arabidopsis seedlings, the unit of replication for biomass weighing was the plate containing 5 seedlings, rather than individual seedlings. Post-tissue collection, fresh weights were recorded using a Mettler Toledo scale, followed by drying the tissues in an oven at 50°C for four days to obtain dry weights. Subsequently, per-plant weights were calculated for further analysis.

To ensure consistent sample replication under both regular and high-temperature conditions, we initiated both experiments simultaneously. Employing the established procedure, plants were grown on MS plates, with the only deviation being the temperature regimen for the high-temperature trial. Specifically, the high temperature was maintained at 28°C during light (16h) exposure and 21°C during dark (8h) periods. Following the same harvesting procedure as the regular temperature experiment, we proceeded to weigh the biomass using the protocol outlined in the preceding paragraph.

### Expression data and GO enrichment of co-expressed genes

We examined published datasets to explore the expression variations of our candidate genes. For organ- and development-specific expression patterns, we used data from Klepikova et al. (2016), and for cell-specific expression, we referred to Shahan et al. (2022). Next, we identified 200 co-expressed genes using the AttedII database (https://atted.jp/) and performed GO enrichment analysis on these genes with PlantRegMap (https://plantregmap.gao-lab.org/go.php), employing default parameters.

### Statistical Analysis

We conducted most of our analyses using R (https://www.r-project.org/) and JMP 16 from SAS (https://www.jmp.com/en_us/home.html), generating statistical outputs and figures. For image editing, we utilized Inkscape (https://inkscape.org/). The physical map of accession distribution was created in JMP using the graph builder. Principal Component Analysis (PCA) and correlation analyses were performed using the REML estimation method. Trait value significance was assessed by the Dunnett test, comparing means from multiple experimental groups (T-DNA lines) to a control group (Col-0) to determine significant differences. For multiple comparisons in single-point experiments, significance was determined by one-way or two-way ANOVA with Tukey’s HSD test (p < 0.05). Each experiment was repeated independently at least twice to ensure consistent results.

## Acknowledgement

We would like to express our gratitude to all members of the Busch lab for their valuable input and ideas throughout the experiments. Special thanks to Joanne Chory for the insightful discussions on biomass accumulation strategies in plants and for her helpful suggestions. We also appreciate Griffin Van Amringe, Lauren Ragel, and Kristine Szeto for their early work on T-DNA screening of TRP candidates. Our thanks go to Mathew Simenc for his assistance in obtaining high-quality single-cell RNA-seq images for the GROOT genes. This research was supported by funding from the Salk Harnessing Plants Initiative to W.B.

## Author Contribution

S.R. and W.B. conceived the study and designed the experiments. S.R., A.P.G., and Y.L.C. conducted the experiments and gathered the phenotypic data. S.R. and L.Z. analyzed the data. W.B. supervised the work and provided funding and resources. S.R. and W.B. wrote the manuscript with contributions from all authors. All authors discussed the results and provided feedback on the manuscript.

