## Supplementary Figure for "Balancing Growth: GROOT Genes Link Plant Biomass and Temperature Adaptation in Arabidopsis"

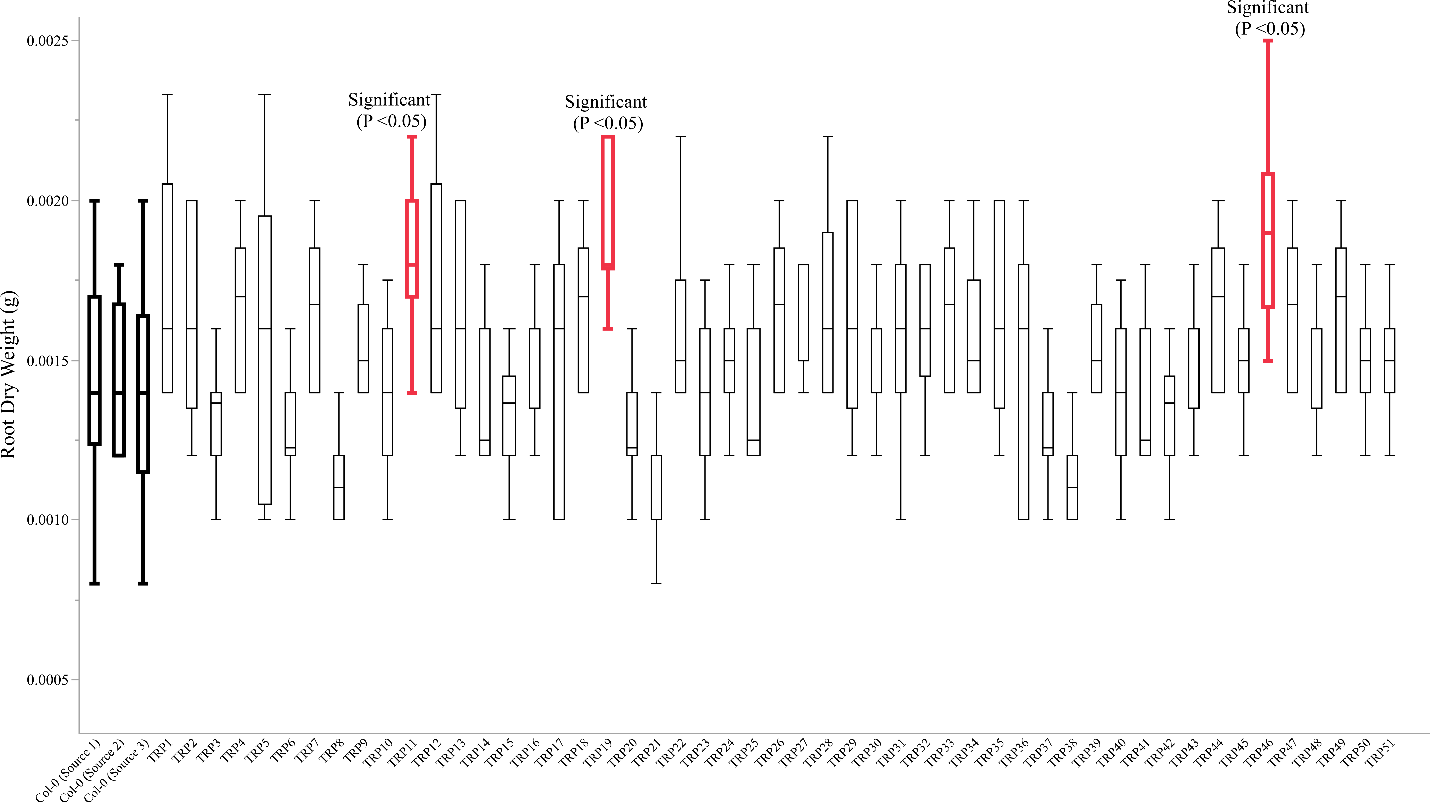

**Supplementary Figure 1**: Root biomass phenotypes were assessed for 51 candidate genes corresponding to the top five GWAS hits. The phenotyping was conducted on day 21 by harvesting the roots, which were then dried at 50°C for four days before weighing. We identified three genes that consistently showed significant root dry weight phenotypes. In this figure, the X-axis represents the names of the T-DNA lines for each of the 51 candidate genes, along with three Col-0 controls collected from distinct sources. The Y-axis represents root dry weight in grams. The data for the Col-0 lines are highlighted in bold black, while the significant genes—TRP11, TRP19, and TRP46, corresponding to the T-DNA lines for *AT3G19440*, *AT3G19590*, and *AT3G19630*, respectively—are shown in red.

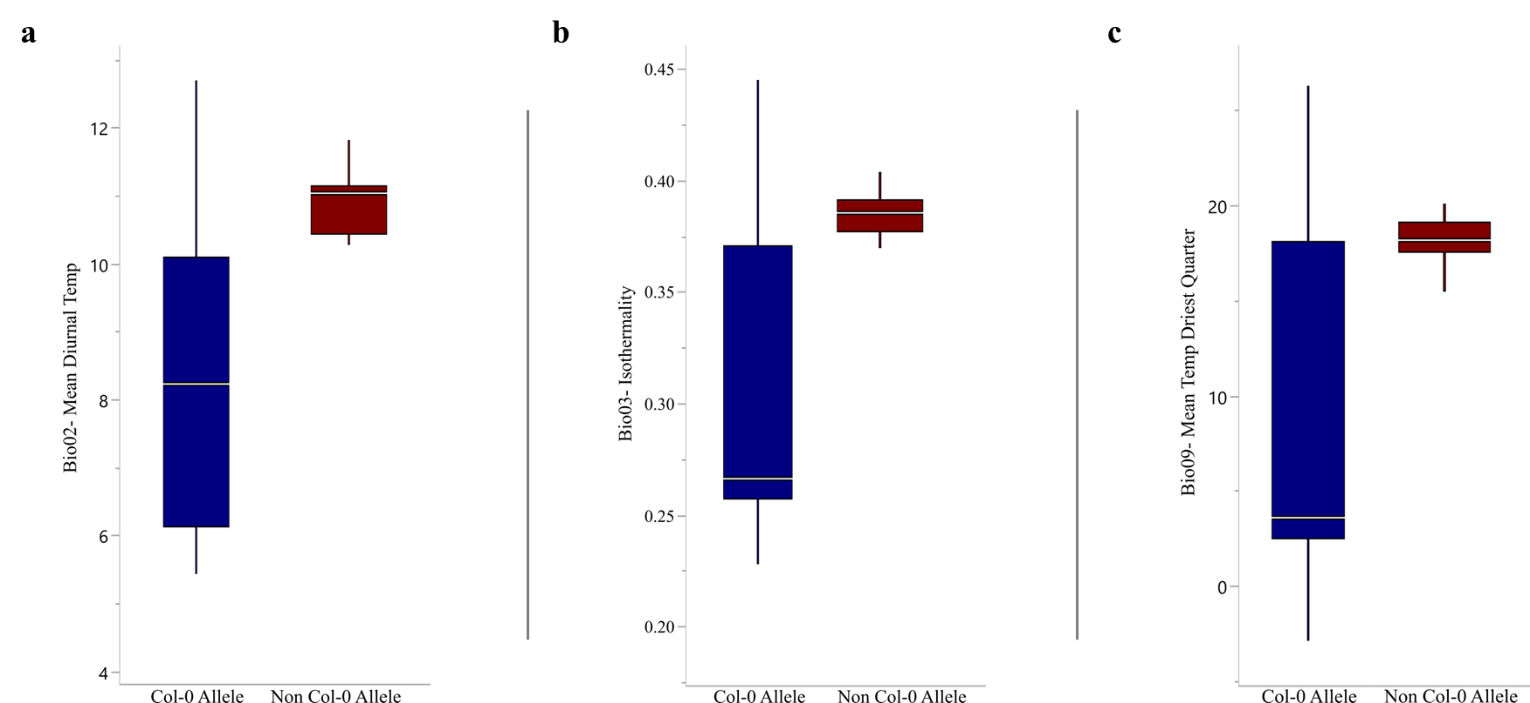

**Supplementary Figure 2**: The relationship between allelic variants at SNP position 6772287 on Chromosome 3 and historical environmental data is shown. For the three significant associations, we observed that accessions containing the non-Col-0 allele consistently had higher values, suggesting they experience greater temperature fluctuations compared to accessions with the Col-0 allele. In all three boxplots, the X-axis represents the allelic variants of accessions for the SNP at position 6772287 on chromosome 3, while the Y-axis denotes the values of specific bioclimatic variables. Accessions with the Col-0 allele are shown in blue, while those with the non-Col-0 allele are shown in red.
